# Chromosomal-level genome assembly of golden birdwing *Troides aeacus* (Felder & Felder, 1860)

**DOI:** 10.1101/2024.01.13.575334

**Authors:** Hong Kong Biodiversity Genomics Consortium, Jerome H.L. Hui, Ting Fung Chan, Leo L. Chan, Siu Gin Cheung, Chi Chiu Cheang, James K.H. Fang, Juan D. Gaitan-Espitia, Stanley C.K. Lau, Yik Hei Sung, Chris K.C. Wong, Kevin Y.L. Yip, Yingying Wei, Wai Lok So, Wenyan Nong, Hydrogen S.F. Pun, Wing Kwong Yau, Colleen Y.L. Chiu, Sammi S.S. Chan, Kacy K.L. Man, Ho Yin Yip

## Abstract

*Troides aeacus*, the golden birdwing (Lepidoptera, Papilionidae) is a large swallowtail butterfly widely distributed in Asia. Despite its occurrence, *T. aeacus* has been assigned as a major protective species in many places given the loss of their native habitats under urbanisation and anthropogenic activities. Nevertheless, the lack of its genomic resources hinders our understanding of their biology, diversity, as well as carrying out conservation measures based on genetic information or markers. Here, we report the first chromosomal-level genome assembly of *T. aeacus* using a combination of PacBio SMRT and Omni-C scaffolding technologies. The assembled genome (351 Mb) contains 98.94% of the sequences anchored to 30 pseudo-molecules. The genome assembly also has high sequence continuity with scaffold length N50 = 12.2 Mb. A total of 28,749 protein-coding genes were predicted, and high BUSCO score completeness (98.9% of BUSCO metazoa_odb10 genes) was also revealed. This high-quality genome offers a new and significant resource for understanding the swallowtail butterfly biology, as well as carrying out conservation measures of this ecologically important lepidopteran species.

## Introduction

The golden birdwing butterfly *Troides aeacus* (Figure 1A) is a swallowtail butterfly that are widely distributed in Asia, including Bangladesh, Burma, Cambodia, China, India, Laos, Malaysia, Nepal, Thailand, and Vietnam (Böhm et al 2020). The species is generally large in size with wingspan reaching ∽15 cm, and has iconic black forewings and golden-yellow hindwings carved with grey stripes and black spots (Wu et al 2010; Li et al 2010). Due to its attractive appearances, it has also been vastly collected and traded in curio markets (Collins and Morris 1985, New and Collins 1991; Wu et al 2010).

**Figure 1.**
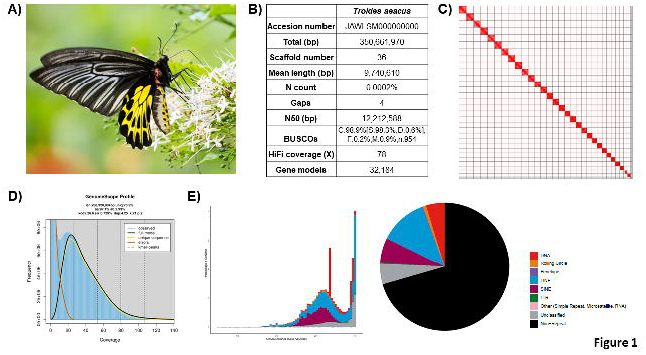
Genomic information of *Troides aeacus*. **A)** Photo of *T. aeacus*; **B)** Statistics of the assembled genome; **C)** Omni-C contact map of the assembly visualised using Juicebox (v1.11.08); **D)** Genomescope report with k-mer = 21; **E)** Repetitive elements distribution in the assembled genome.

Similar to other homometabolans, *T. aeacus* exhibits larvae and pupae stages (5 larval instar stages before transforming into green girdled pupal stage)(Li et al 2010). The larvae are generally dependent on host plants in the Aristolochiaceae, especially of the genus *Aristolochia* that can be commonly found in Asia (Li et al 2010; Wu et al 2010; Böhm et al 2020; Chen et al 2013). After emergence, the adults will feed and live around nectaring flowers such as those in the genus *Hibiscus, Ixora, Lantana, Mussaenda*, and *Spathodea* (Cotton and Racheli 2006; Böhm et al 2020). Anthropogenic activities including deforestation, grazing, herbicide application, hunting, land reclamation, mine exploitation, and trading all have been suggested to pose threats to *T. aeacus* (Böhm et al 2020; Li et al 2010; Khanal 2022). In certain places such as Hong Kong, *T. aeacus* has also been suggested to be under protection and restoration of its lost; and in Taiwan, the trade of endemic subspecies such as *T. aeacus* is protected by the Convention on International Trade in Endangered Species of Wild Fauna and Flora (CITES) (Fellowes et al 2002).

## Context

To date, genomic resource in the genus *Troides* are confined to *T. helena* (He et al 2022) and *T. oblongomaculatus* (Reboud et al 2023). In light of the high conservation value of *T. aeacus* and its phylogenetic position to understand the diversification of butterflies (Condamine et al 2015), it has been chosen as one of the species for genome sequencing by the Hong Kong Biodiversity Genomics Consortium (a.k.a. EarthBioGenome Project Hong Kong), which is formed by investigators from 8 publicly funded universities in Hong Kong. Here, we report a chromosomal-level genome assembly of the golden birdwing *T. aeacus*.

## Methods

### Sample collection and species identification

A pupa of the golden birdwing *Troides aeacus* was obtained in August 2022. The pupa was snap-frozen in liquid nitrogen upon collection. The frozen pupa was further grinded into fine powder and stored at -80 °C freezer until DNA isolation. A portion of the powder was used for species molecular identification with QIAamp DNA Mini Kit (Qiagen Cat. 51306), following the provided protocol. The DNA was then used as a template for conventional PCR with the following protocol: an initial denaturation step at 95 °C for 3 minutes; followed by 36 amplification cycles of 30 seconds for denaturation at 95 °C; 30 seconds for primer annealing at 55 °C and 1 minute for extension at 72 °C; and a final extension step at 72 °C for 3 minutes. The reaction mixture included PCR buffer, DNA template, 2 mM dNTP, 1.5 mM MgCl_2_, 0.4 mM of each forward and reverse primers (LCO1490: 5’-GGTCAACAAATCATAAAGATATTGG-3’, HCO2198: 5’-TAAACTTCAGGGTGACCAAAAAATCA-3’) (Folmer et al 1994), and *Taq* DNA polymerase. The PCR was performed on a T100™ thermal cycler (Bio-Rad, USA).

### Isolation of high molecular weight genomic DNA

High molecular weight genomic DNA was isolated with the remaining stored powder using the Qiagen MagAttract HMW kit (Qiagen Cat. No. 67563) following the protocol. In summary, 1 g of sample powder was placed in microcentrifuge tube with 200 μl 1X phosphate-buffered saline (PBS), RNase A, Proteinase K, and Buffer AL. The mixture was incubated at room temperature (22-25 °C) for 2.5 hours, until the tissue was completely disintegrated. The sample was eventually eluted with 120 μl of elution buffer (PacBio Ref. No. 101-633-500) and stored at 4 °C. In order to keep the integrity of the DNA, wide-bore pipette tips were used in any transfer of DNA during the process. The sample was then proceeded to quality control by quantifying with the Qubit^®^ Fluorometer and Qubit^™^ dsDNA HS and BR Assay Kits (Invitrogen^™^ Cat. No. Q32851). An overnight pulse-field gel electrophoresis was performed to estimate the size of the isolated DNA, with three markers (λ-Hind III digest; Takara Cat. No. 3403, DL15,000 DNA Marker; Takara Cat. No. 3582A and CHEF DNA Size Standard-8-48 kb Ladder; Cat. No. 170-3707). Besides, the sample purity was examined by the NanoDrop^™^ One/OneC Microvolume UV-Vis Spectrophotometer (with A260/A280: ∽1.8 and A260/A230: >2.0 as a standard threshold).

### DNA shearing, PacBio library preparation and sequencing

120 μl of DNA sample of 10 μg DNA was transferred to a g-tube (Covaris Part No. 520079). The tube was then allowed to proceed 6 passes of centrifugation with 2,000 x g of 2 minutes each. The resultant DNA was saved in a 2 mL DNA LoBind^®^ Tube (Eppendorf Cat. No. 022431048) at 4 °C until library preparation. The molecular weight of the isolated DNA was examined by overnight pulse-field gel electrophoresis. The electrophoresis profile was set as follow: 5K as the lower end and 100K as the higher end for the designated molecular weight; Gradient = 6.0V/cm; Run time = 15 h:16 min; included angle = 120 °; Int. Sw. Tm = 22 s; Fin. Sw. Tm = 0.53 s; Ramping factor: a = Linear. The gel was run in 1.0% PFC agarose in 0.5X TBE buffer at 14 °C.

A SMRTbell library was made using the SMRTbell® prep kit 3.0 (PacBio Ref. No. 102-141-700), following the provided protocol. In summary, single-stranded overhangs of the genomic DNA were removed, and the DNA is repaired from physical damage during shearing. Subsequently, both DNA ends were tailed with an A-overhang and ligation of T-overhang SMRTbell adapters was performed at 20 °C for 30 minutes. The SMRTbell library was then purified with SMRTbell^®^ cleanup beads (PacBio Ref. No. 102158-300). The size and concentration of the library were assessed with the pulse-field gel electrophoresis and the Qubit^®^ Fluorometer and Qubit™ dsDNA HS and BR Assay Kits (Invitrogen^™^ Cat. No. Q32851), respectively. A subsequent nuclease treatment step was carried out to remove non-SMRTbell structures in the library. A final size-selection step was performed to remove small DNA fragments in the library with 35% AMPure PB beads. The Sequel^®^ II binding kit 3.2 (PacBio Ref. No. 102-194-100) was used for final preparation. In short, Sequel II primer 3.2 and Sequel II DNA polymerase 2.2 were annealed and bound to the SMRTbell library, respectively. The library was loaded afterwards at an on-plate concentration of 50-90 pM using the diffusion loading mode. The sequencing was conducted on the Sequel IIe System with an internal control provided in the kit. The sequencing was performed in 30-hour movies, with 120 min pre-extension, connected to the software SMRT Link v11.0 (PacBio). HiFi reads are eventually generated and collected for further analysis. One SMRT cell was used for this sequencing (Supplementary Information 1).

### Omni-C library preparation and sequencing

An Omni-C library was made using the Dovetail® Omni-C® Library Preparation Kit (Dovetail Cat. No. 21005) according to the provided protocol. In summary, 80 mg of frozen powered tissue sample was placed in a microcentrifuge tube with 1 mL 1X PBS and formaldehyde. The fixed DNA was digested with endonuclease DNase I. Afterwards, the concentration and size of the digested sample was examined by the Qubit® Fluorometer and Qubit™ dsDNA HS and BR Assay Kits (Invitrogen™ Cat. No. Q32851), and the TapeStation D5000 HS ScreenTape, respectively. Both DNA ends were the polished and ligation of biotinylated bridge adaptor was conducted at 22 °C for 30 minutes. Subsequent proximity ligation between crosslinked DNA was performed at 22 °C for 1 hour. After ligation, the DNA was reverse crosslinked, and purified with SPRIselect™ Beads (Beckman Coulter Product No. B23317) to remove the biotin that was not internal to the ligated fragments. The Dovetail™ Library Module for Illumina (Dovetail Cat. No. 21004) was used for end repair and adapter ligation. The DNA was tailed with an A-overhang, which allowed Illumina-compatible adapters to ligate to the DNA fragments at 20 °C for 15 minutes. The Omni-C library was then sheared into fragments with USER Enzyme Mix and further purified with SPRIselect™ Beads. Isolation of DNA fragments with internal biotin were performed with Streptavidin Beads. Universal and Index PCR Primers from the Dovetail™ Primer Set for Illumina (Dovetail Cat. No. 25005) were used to amplify the constructed library. Size selection was carried out with SPRIselect™ Beads targeting fragments ranging between 350 bp and 1000 bp. Finally, the concentration and fragment size of the sequencing library was examined with the Qubit® Fluorometer and Qubit™ dsDNA HS and BR Assay Kits, and the TapeStation D5000 HS ScreenTape, respectively. The resultant library was sequenced on the Illumina HiSeq-PE150 platform (Supplementary Information 1).

### Genome assembly and gene model prediction

*De novo* genome assembly was performed using Hifiasm (Cheng et al 2021). Haplotypic duplications were identified and removed using purge_dups based on the depth of HiFi reads (Guan et al 2020). Proximity ligation data from the Omni-C library were used to scaffold genome assembly by YaHS (Zhou et al., 2022). Transposable elements (TEs) were annotated as previously described (Baril et al 2022), using the automated Earl Grey TE annotation pipeline (version 1.2, https://github.com/TobyBaril/EarlGrey). Gene models were predicted from the soft-masked genome assembled by funannotate (Palmer & Stajich, 2020) using predict with the parameters “--protein_evidence uniprot_sprot.fasta --genemark_mode ES --optimize_augustus --busco_db metazoa --organism other --max_intronlen 350000” and other default parameters.

## Results and discussion

### Genome assembly of T. aeacus

A total of 27 Gb of HiFi bases were yielded with an average HiFi read length of 9,688 bp with 78X coverage (Supplementary Information 1). After incorporating with 21.7 Gb Omni-C data, the resulting genome assembly was 350.66 Mb in size with 36 scaffolds, of which 30 scaffolds are of chromosome length (Figure 1B-C; Table 1-2). The genome has a high contiguity with a scaffold N50 value of 12.21 Mb and high completeness with the complete BUSCO estimation to be 98.9% (metazoa_odb10)(Figure 1B, Table 1). While the genome size estimation was about 268.3 Mb with 2.93% nucleotide heterozygosity rate (Figure 1D; Supplementary Information 2), the assembled *Troides aeacus* genome has a genome size similar to other swallowtail butterfly genomes, including *Troides Helena* (∽330 Mb) (He et al 2022) and *Troides oblongomaculatus* (∽348 Mb) (Reboud et al 2023). In addition, 43 telomeres were found in 25 scaffolds of the assembly genome (Table 3). Furthermore, 32,183 gene models were predicted with a BUSCO score of 86.5%.

### Repeat content

A total repetitive content of 29.50% was identified in the assembled genome, including 5.16% unclassified elements (Figure 1E; Table 4). Among the known repeats, LINE is the most abundant (12.01%), followed by SINE retrotransposons (6.38%) and DNA transposons (4.71%), whereas Rolling Circle, LTR, Penelope and other are present in low proportions (Rolling Circle: 0.78%, LTR: 0.26%, Penelope: 0.20%, other: 0.02%).

## Conclusion and reuse potential

This study presents the first chromosomal-level genome assembly of the golden birdwing *Troides aeacus*, which is a useful and precious resource for further phylogenomic studies of birdwing butterfly species in light of species diversification and conservation.

## Data validation and quality control

During DNA extraction and PacBio library preparation, the samples were subjected to quality control with NanoDrop^™^ One/OneC Microvolume UV-Vis Spectrophotometer, Qubit^®^ Fluorometer, and overnight pulse-field gel electrophoresis. The Omni-C library was inspected by Qubit^®^ Fluorometer and TapeStation D5000 HS ScreenTape.

Regarding the genome assembly, the Hifiasm output was blast to the NT database and the resultant output was used as the input for BlobTools (v1.1.1) (Laetsch & Blaxter 2017). Scaffolds that were identified as possible contamination were removed from the assembly manually (Supplementary Information 3). A statistical kmer-based approach was applied to estimate the heterozygosity of the assembled genome heterozygosity The repeat content and the corresponding sizes were analysed using Jellyfish (Marçais & Kingsford 2011) and GenomeScope (Ranallo-Benavidez et al 2020) (Figure 1D; Supplementary Information 2). Furthermore, telomeric repeats were inspected by FindTelomeres (https://github.com/JanaSperschneider/FindTelomeres). Benchmarking Universal Single-Copy Orthologs (BUSCO, v5.5.0) (Manni et al., 2021) was used to assess the completeness of the genome assembly and gene annotation with metazoan dataset (metazoa_odb10).

## Data availability

The final genome assembly was submitted to NCBI under accession numbers GCA_033220335.2 (https://www.ncbi.nlm.nih.gov/datasets/genome/GCA_033220335.2). The raw reads yielded from this study were deposited to the NCBI database under the BioProject accession number PRJNA973839. The genome annotation files were deposited in the figshare (https://figshare.com/s/811f9d4158e592e9d209).

## Author’s contributions

JHLH, TFC, LLC, SGC, CCC, JKHF, JDG, SCKL, YHS, CKCW, KYLY and YW conceived and supervised the study; WLS carried out DNA extraction, library preparation and sequencing; WN performed genome assembly and gene model prediction; HSFP, WKY, CYLC, SSSC and KKLM and HYY collected and maintained the butterfly samples. All authors approved the final version of the manuscript.

## Acknowledgements

The authors thank Sean Law for discussion.

## Figure legends

**Table 1.** Details of genome assembly statistics.

**Table 2.** Information of 30 chromosomal-length scaffolds.

**Table 3.** Summary of telomeric repeats found in 25 scaffolds.

**Table 4.** Summary of repetitive elements in the genome.

**Supplementary Information 1**. Summary of genome sequencing data.

**Supplementary Information 2**. GenomeScope result summary (k-mer = 21).

**Supplementary Information 3**. Genome assembly QC and contaminant detection.

