## Supplementary Information 1 for "Chromosomal-level genome assembly of golden birdwing *Troides aeacus* (Felder & Felder, 1860)": EBPHK Butterfly Supplementary Information.docx

**Supplementary Information 1.** Summary of genome sequencing data.

| **Library** | **Reads** | **Bases** | **Coverage(X)** | **Accession** |
| --- | --- | --- | --- | --- |
| PacBio HiFi | 2,805,656 | 27,181,071,888 | 78 | SRR24631717 |
| Omni-C | 144,777,842 | 21,716,676,300 | 62 | SRR26815782 |

**Supplementary Information 2.** GenomeScope result summary (k-mer = 21).

| Property | Min | Max |
| --- | --- | --- |
| Homozygous (aa) | 97.04% | 97.11% |
| Heterozygous (ab) | 2.89% | 2.96% |
| Genome Haploid Length (bp) | 265,767,007 | 268,320,884 |
| Genome Repeat Length (bp) | 69,335,920 | 70,002,201 |
| Genome Unique Length (bp) | 196,431,086 | 198,318,683 |
| Model Fit | 78.57% | 99.17% |
| Read Error Rate | 0.73% | 0.73% |

**Supplementary Information 3.** Genome assembly QC and contaminant detection.

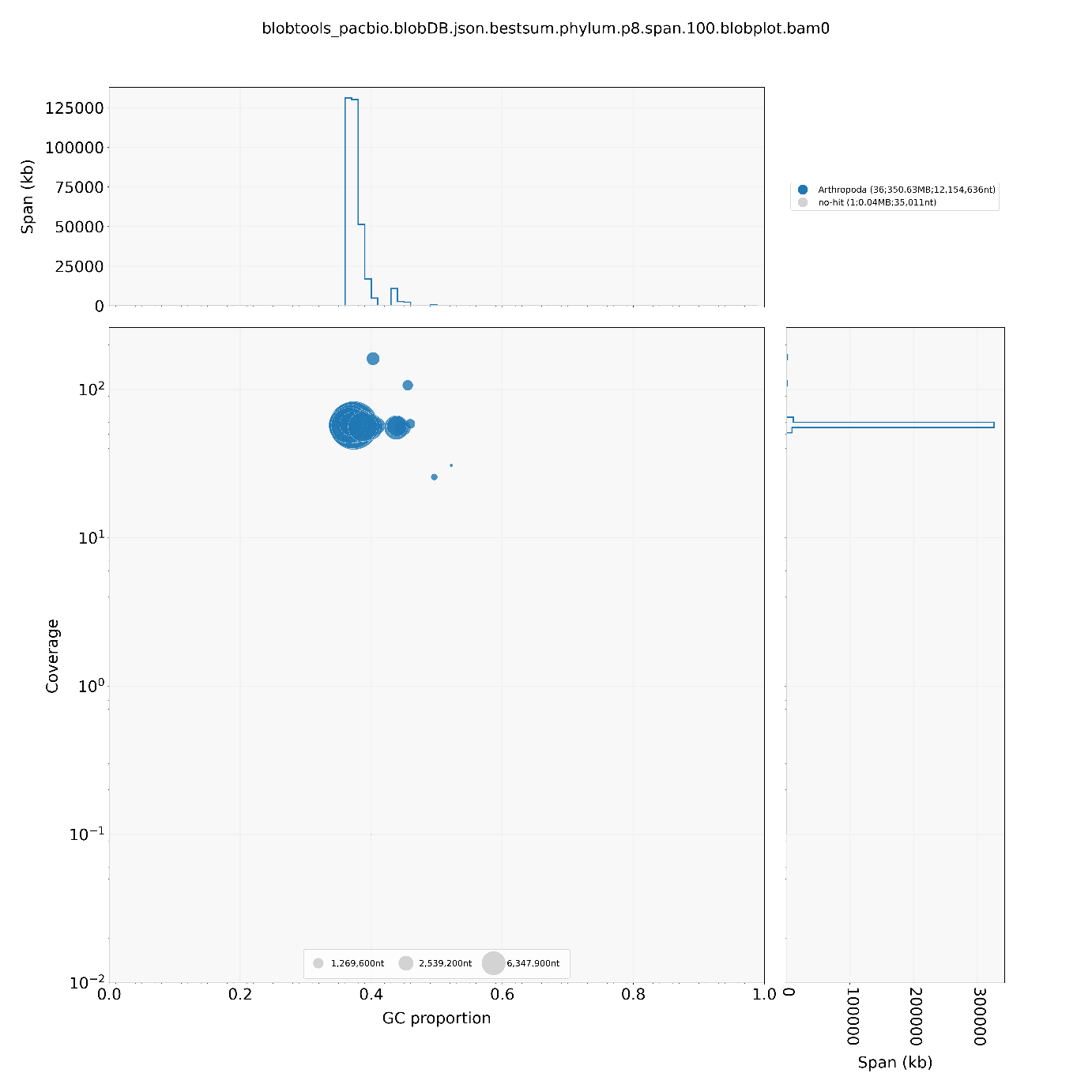

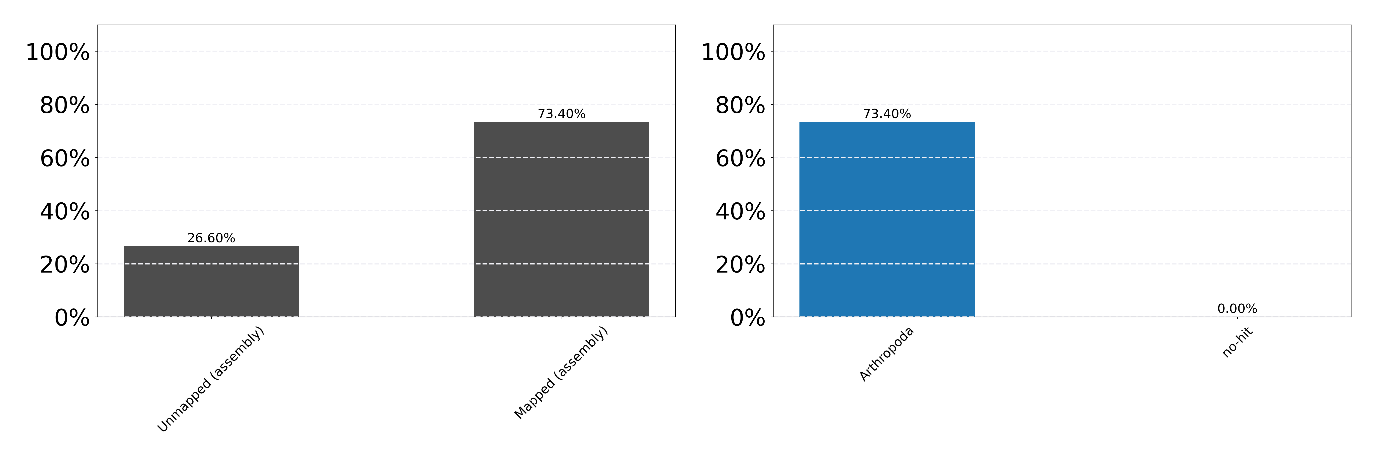
